# Isobaric matching between runs and novel PSM-level normalization in MaxQuant strongly improve reporter ion-based quantification

**DOI:** 10.1101/2020.03.30.015487

**Authors:** Sung-Huan Yu, Pelagia Kiriakidou, Jürgen Cox

## Abstract

Isobaric labeling has the promise of combining high sample multiplexing with precise quantification. However, normalization issues and the missing value problem of complete n-plexes hamper quantification across more than one n-plex. Here we introduce two novel algorithms implemented in MaxQuant that substantially improve the data analysis with multiple n-plexes. First, isobaric matching between runs (IMBR) makes use of the three-dimensional MS1 features to transfer identifications from identified to unidentified MS/MS spectra between LC-MS runs in order to utilize reporter ion intensities in unidentified spectra for quantification. On typical datasets, we observe a significant gain in quantifiable n-plexes. Second, we introduce a novel PSM-level normalization, applicable to data with and without common reference channel. It is a weighted median-based method, in which the weights reflect the number of ions that were used for fragmentation. On a typical dataset, we observe complete removal of batch effects and dominance of the biological sample grouping after normalization. Furthermore, we provide many novel processing and normalization options in Perseus, the companion software for the downstream analysis of quantitative proteomics results. All novel tools and algorithms are available with the regular MaxQuant and Perseus releases, which are downloadable at http://maxquant.org.

---

Mass spectrometry (MS) has revolutionized the way researchers can monitor protein concentration changes on a proteome-wide scale. Several techniques have been established for quantifying relative amounts of proteins or peptides between related samples as, for instance, stable isotope labeling on the level of first-stage MS (MS1) spectra^1–4^, label-free quantification^5,6^ (LFQ) and isobaric labeling^7–9^. The latter has the advantage of allowing for relatively high sample multiplexing and often is done in the form of tandem mass tags^7^ (TMT) or isobaric tags for relative and absolute quantitation^7,10^ (iTRAQ). Isobaric labeling can substantially improve on a genuine issue for shotgun proteomics, which is the missing value problem. In unlabeled samples, values for quantification can be missing due to several reasons, as for instance low abundance of the protein or lack of identification of peptides^11,12^. It has been observed that in label-free samples the fraction of proteins containing one or more missing value can dominate the proteome data^13^. Isobaric labeling improves the situation to some extent, since within an n-plex the chances of getting missing values are strongly reduced. However, the missingness of complete n-plexes occurs with the same likelihood as single values are missing in label-free data.

MaxQuant is one of the most widely used platforms for analyzing shotgun proteomics data^14–17^. In order to recover features for quantification, beyond those that were directly identified by fragmentation spectra, several methods have been developed and integrated into the MaxQuant software in the past. Match between runs^18,19^ (MBR) is one of the methods to decrease the number of missing values in label-free and MS1-level labeled data. It can transfer peptide identifications from an LC-MS run, in which the peptide is identified by MS/MS, to another LC-MS run, in which the same peptide exists as an MS1 feature, but was not identified, either because no fragmentation spectrum was recorded for this MS1 feature, or because the recorded fragmentation spectrum was not identified by the peptide search engine^20^. Required for this transfer of identifications between similar samples to happen with low rates of false positives are the high mass accuracy obtained after nonlinear mass recalibration and comparable retention times after retention time alignment in MaxQuant. ‘Re-quantify’ provides another method to increase the coverage of protein quantification for the case of MS1-level labeling. An isotope pattern that has not been paired with any labeling partners can, after it has been identified, be restored for quantification of ratios by integrating the mass spectrometric signal at the expected position in m/z and retention time coordinates in the same LC-MS run^20^. Although these options can improve the protein quantification for label-free or MS1-labeling experiments, a method of recovering values for isobaric labeling still needed to be developed in MaxQuant.

Besides procedures for recovering missing values, normalization is another important step in isobaric labeling-based quantification. For removing batch effects, several useful normalization methods have recently been developed^21–27^. Almost all of the methods are applying corrections at the level of protein quantification. An exception is the ‘compositional proteomics’ strategy^28^ which removes effects due to constraints imposed on the sum over channels for each PSM. We propose a new and straightforward PSM-level normalization method based on the weighted median of ratios. For MS1-level labeling it is advantageous to define the protein ratio as the median of the peptide feature ratios, as is done in MaxQuant. For MS1 signals, which are not affected by co-fragmentation, the gain in robustness resulting from applying a median approach to the ratios outweighs the lack of weighing ratios by their signal strength. This is not the case for isobaric labeling signals, which often show inferior results if the protein ratio is taken as the median of PSM ratios. Rather simplistic methods that are not robust in the sense of the median, but weighted by signal intensity are better suited here, since they reduce the influence of co-fragmentation. The novel normalization method that we are proposing combines the strength of both approaches, the robustness of the median and the weighting of PSMs by signal strength.

In this manuscript, we present a novel isobaric match between runs (IMBR) and a PSM-level normalization method that were built into MaxQuant. They can significantly increase the number of peptide features that are available for quantification and efficiently remove the batch effects. Moreover, numerous plug-ins of Perseus^29,30^, which is a powerful platform for the downstream analysis of data generated with MaxQuant or other platforms, have been developed for processing isobaric labeling datasets. All the software is available for download at http://maxquant.org.

## ■ EXPERIMENTAL SECTION (1080/1500)

### Datasets

For the evaluation of our newly developed methods, we use well-established datasets that were submitted to open access community databases. We downloaded isobaric labeling datasets of two types: those with a reference channel, in which one of the channels carries the signal of the mixture of samples suitable for forming ratios to the other channels that carry the actual samples, and those without a reference channel. As an example for the latter we obtained data from Bailey *et al*.^31^. It is an 8-plex dataset consisting of eight organs (kidney, lung, heart, muscle, liver, cerebrum, cerebellum, spleen) harvested from four different mice. After tryptic digestion, the peptides of each organ were labeled with a TMT 8-plex in a randomized design and applied to two different data acquisitions - data dependent acquisition (DDA) and Intelligent data acquisition (IDA). The dataset was obtained from the Chorus database (http://chorusproject.org/; 298: Elution Order Algorithm). Moreover, we downloaded a TMT 10-plex labeling dataset containing a reference channel acquired by Lereim *et al*.^32^. It consists of brain tissues from WT mice and *Peli1* knock-out mice. *Peli1* functions as a regulator of the immune response during experimental autoimmune encephalomyelitis (EAE). Each type of mice contains three samples based on the number of days post EAE infection: 0, 10 and 20. The samples are randomly assigned to two TMT 10-plex sets, and the TMT131 is for the pooled samples consisting of a mixture of same amounts of all samples. The dataset is available in PRIDE^33^ (PXD003710).

### Data processing

For both datasets we used mouse UniProt sequences (UP000000589, reviewed at 24-07-2018, 16992 proteins). All searches were performed with oxidation of methionine and protein N-terminal acetylation as variable modifications and cysteine carbamidomethylation as fixed modification. Trypsin was selected as protease allowing for up to two missed cleavages, and the peptide mass was limited to a maximum of 4600Da. The initial mass tolerance was 20 ppm for precursor ions and 20 ppm for fragment ions. PSM and protein FDRs were both applied at 1%. In general, values of parameters in MaxQuant have not been changed from their default values unless explicitly stated.

### Isobaric match between runs

Matching between runs of MS1 features is performed exactly as it was done before for the quantification of unlabeled samples (Figure 1). Prior to matching of MS1 features, their masses are recalibrated and their retention times are aligned in MaxQuant. The yellow MS1 isotope pattern in Figure 1 appears in both LS-MS runs. It has been identified by an MS/MS spectrum in the left run. In the right run, the same MS1 feature appears, however with an MS/MS spectrum that did not lead to the identification of the peptide due to poor coverage of the y and b ion series. This MS/MS spectrum does however contain reporter ion intensities, which is then used for quantification.

**Figure 1.**
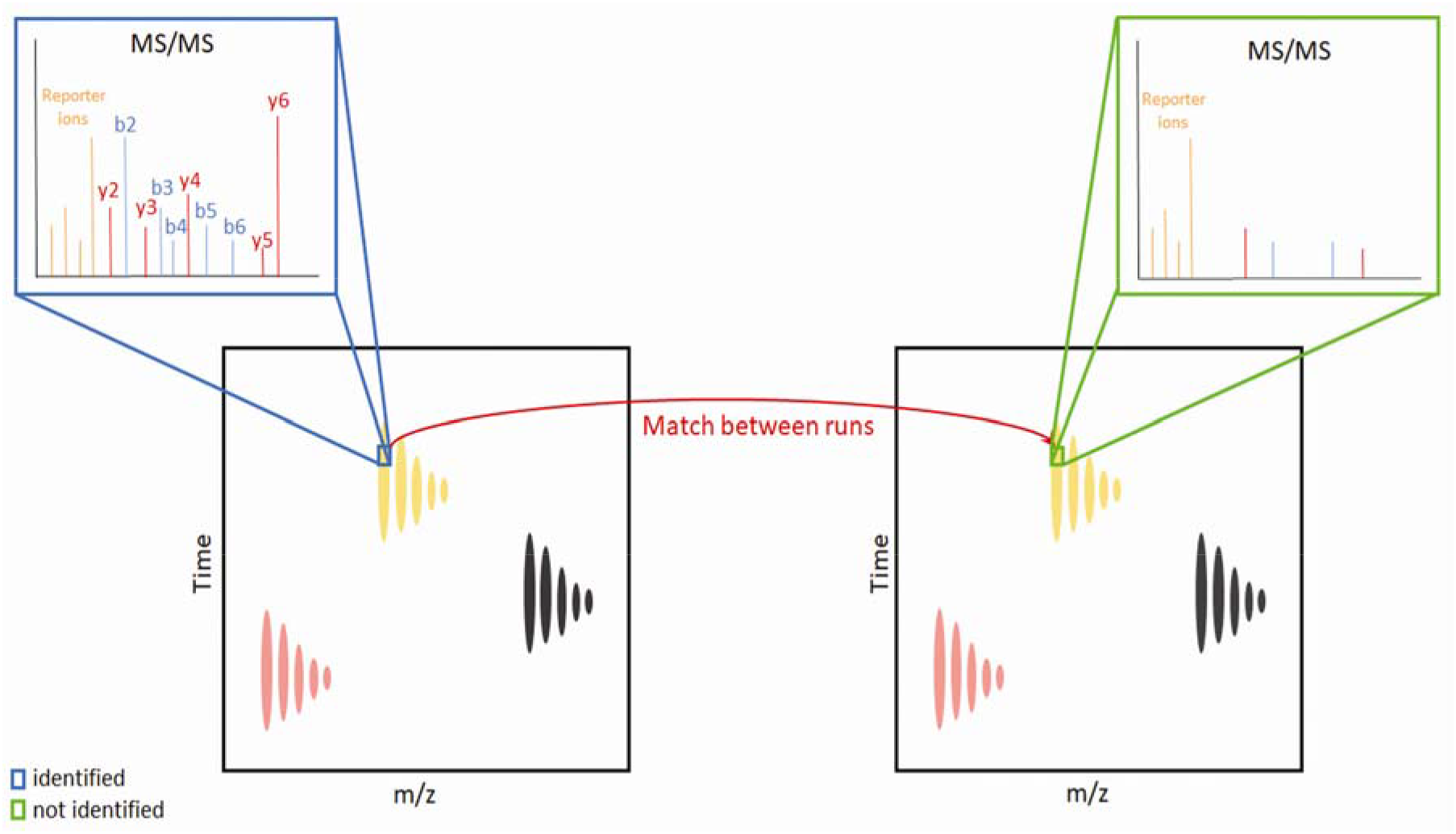
Schematics of IMBR. The reporter ions in the unidentified MS/MS spectrum can be used for quantification based on the matching of MS1-level 3D isotope patterns.

### PSM-level weighted ratio normalization

In order to remove batch effects, we developed a novel method for normalizing reporter ion intensities in isobaric labeling datasets at the PSM-level and integrated it into the MaxQuant software. First, for each protein, the quantifiable PSMs are retrieved. These follow in the first instance the same rules as for label-free or MS1-level labeling quantification. For instance, if the protein quantification should be based only on protein group-level unique peptide sequences, then these are selected. On top of this, MS2-level labeling specific filters can be applied, as for instance based on precursor ion fraction (PIF) or base peak ratio. The resulting set of filtered PSMs then is subjected to a weighted median calculation of reporter ion intensities. This is done for raw intensities as well as intensities corrected for channel mixing, which are both reported in the output tables. The weighted median of the ratios *x*_*1*_, *x*_*2*_, …, *x_*n*_* for a particular isobaric labeling channel to the reference channel over *n* valid PSMs matching to the protein group is then taken. For the weighted median calculation, we assume that the ratios *x*_*i*_ are sorted in ascending order and that the positive weights *w*_*1*_, *w*_*2*_, …, *w*_*n*_ sum up to one. The weighted median is the ratio *x*_*k*_ with the index *k* satisfying

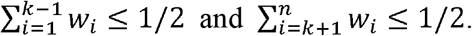

In case that any of the inequalities is an equality, appropriate generalizations apply^34^. The weights *w*_*i*_ are the product of the precursor ion intensity exactly at the retention time at which the MS/MS has been recorded times the fill time of the MS/MS spectrum. This is supposed to be proportional to the number of ions that are used for fragmentation. The weights are then exponentiated with a constant which can be set by the user (‘Isobaric weight exponent’ parameter in the graphical user interface), and whose optimal value is determined as follows.

### Analysis workflow in MaxQuant and Perseus

For running isobaric labeling data in MaxQuant, first, the reference channel(s) need to be assigned after loading the raw data. Multiple reference channels per n-plex are supported, in which case the sum of signals over the reference channels is used for normalization. If the dataset does not contain reference channels, all channels have to be assigned as reference channels and their total signal sum is taken in that case. Secondly, the ‘Reporter ion MS2’ has to be selected as ‘Type’ under ‘Group-specific parameters’, and the suitable set of isobaric labels has to be chosen. Furthermore, if isobaric match between runs should be applied, the ‘Match between runs’ option in ‘Identification’ under ‘Global parameters’ needs to be turned on. In addition, ‘Normalization’ needs to be specified as ‘Ratio to reference channel’, which is normalizing the data without weight or ‘Weighted ratio to reference channel’. If ‘Weighted ration to reference channel’ is selected, the weight can be defined in ‘Isobaric weight exponent’ on the ‘Misc.’ page. The parameters of newly developed methods are shown in Figure S1.

For analysis of the output tables of MaxQuant in Perseus, protein groups which are known contaminants, only identified by site, reverse, or of low quality regarding missing values (more than 30% values are missing in total) are removed. Moreover, the annotations of cell types were generated and the imputation of remaining missing values was performed based on sampling from a normal distribution (width is 0.3 and down-shift is 1.8) for UMAP or other downstream analysis. Additionally, the columns of reference channels were excluded from the analysis. The workflow is presented in Figure S2. For the details of the newly developed plugins for isobaric labeling data analysis and dimensionality reduction methods are shown in Figure S3 and S4.

## ■ RESULTS AND DISCUSSION

### Reduction of missing values

To study the influence of IMBR on missing values we performed MaxQuant analyses once without and once with IMBR with all other parameters at default values as described in the Experimental Section on both datasets. An evidence entry corresponds to a 3D MS1 feature that has at least one MS/MS spectrum attached that is used for quantification. On both datasets, we see a consistent increase of evidence entries by approximately 50% in MS1 level and 7-9% in MS2 level through IMBR. In Figure 2, the distribution of peptides found in a certain number of isobaric labeling batches is displayed with and without IMBR for the dataset by Bayley et al. As can be seen, by applying IMBR, the amounts of quantified peptides and proteins found in all samples are 2.5 and 1.5 fold more than without using IMBR. In particular situations for which peptide level quantification is necessary, as for instance in the study of posttranslational modifications like phosphorylation, the increase in quantified n-plexes is particularly substantial.

**Figure 2.**
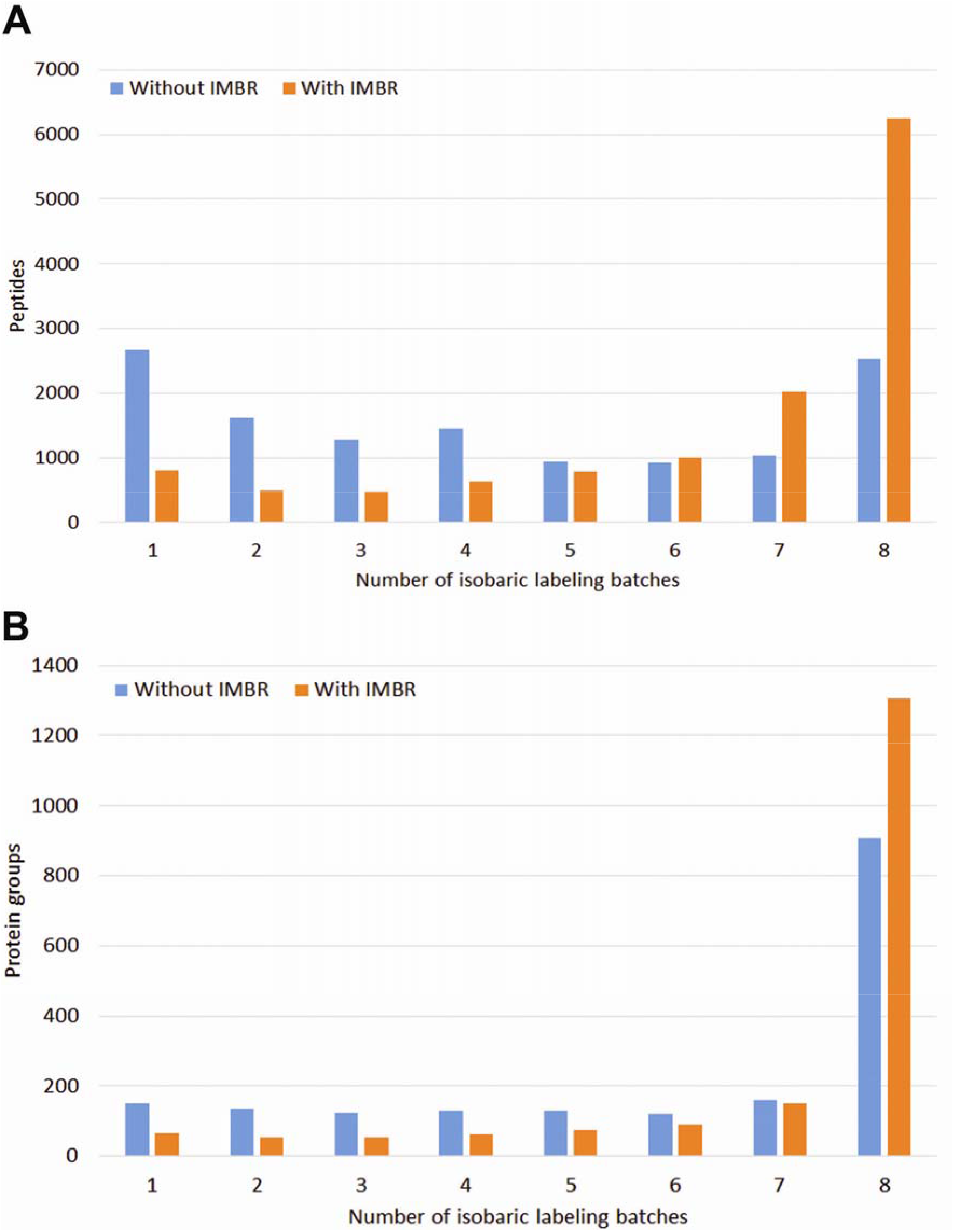
Improvement by IMBR. **A.** For the Bailey et al. dataset it is shown how many peptides are found in *n* out of eight isobaric labeling batches. Blue and orange bars represent results without and with IMBR, respectively. **B.** Same as A. for protein groups.

### Determination of the optimal isobaric weight exponent

For determining the optimal value for the parameter ‘Isobaric weight exponent’, we scan W different values of the weight parameter between 0 and 1 with increment 0.05. Consider a dataset with a biological or technical replicate grouping into *G* groups and quantitative data for *P* proteins (or, more specifically, protein groups). The samples within the replicate groups should be completely randomized over the isobaric labeling batches, as it has been done in the datasets we are analyzing. We want to monitor a measure for the variability within replicate groups in relation to the total variability. For this purpose, we define

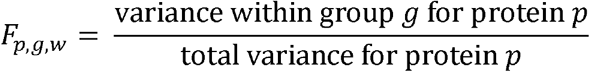

where *w* = *1,…,W* is indexing the different values for the isobaric weight exponent. All variance calculations are performed on logaritmized and imputed values as described above. We take the median of these quantities over all proteins to obtain

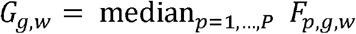

which is a measure for the relative spread of group *g* when using weight parameter *w*. In order to get a balanced contribution from all groups, we rescale the *G*_*g,w*_ within each group over the weight parameter values to a range from 0 to 100,

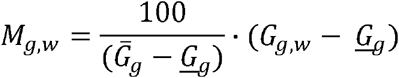

where

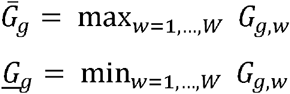

The quantity

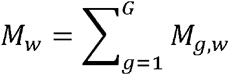

is then optimized over the different values of the isobaric weight exponent. Figure 3 reveals that *M*_*w*_ is minimal if the weight exponent assumes the value 0.75 on the dataset by Bailey et al. Based on these findings, the default value for the isobaric weight exponent is set to 0.75 in the software. Since the optimal value might slightly change between datasets, the user may re-optimize the value for their data and change the value accordingly in MaxQuant.

**Figure 3.**
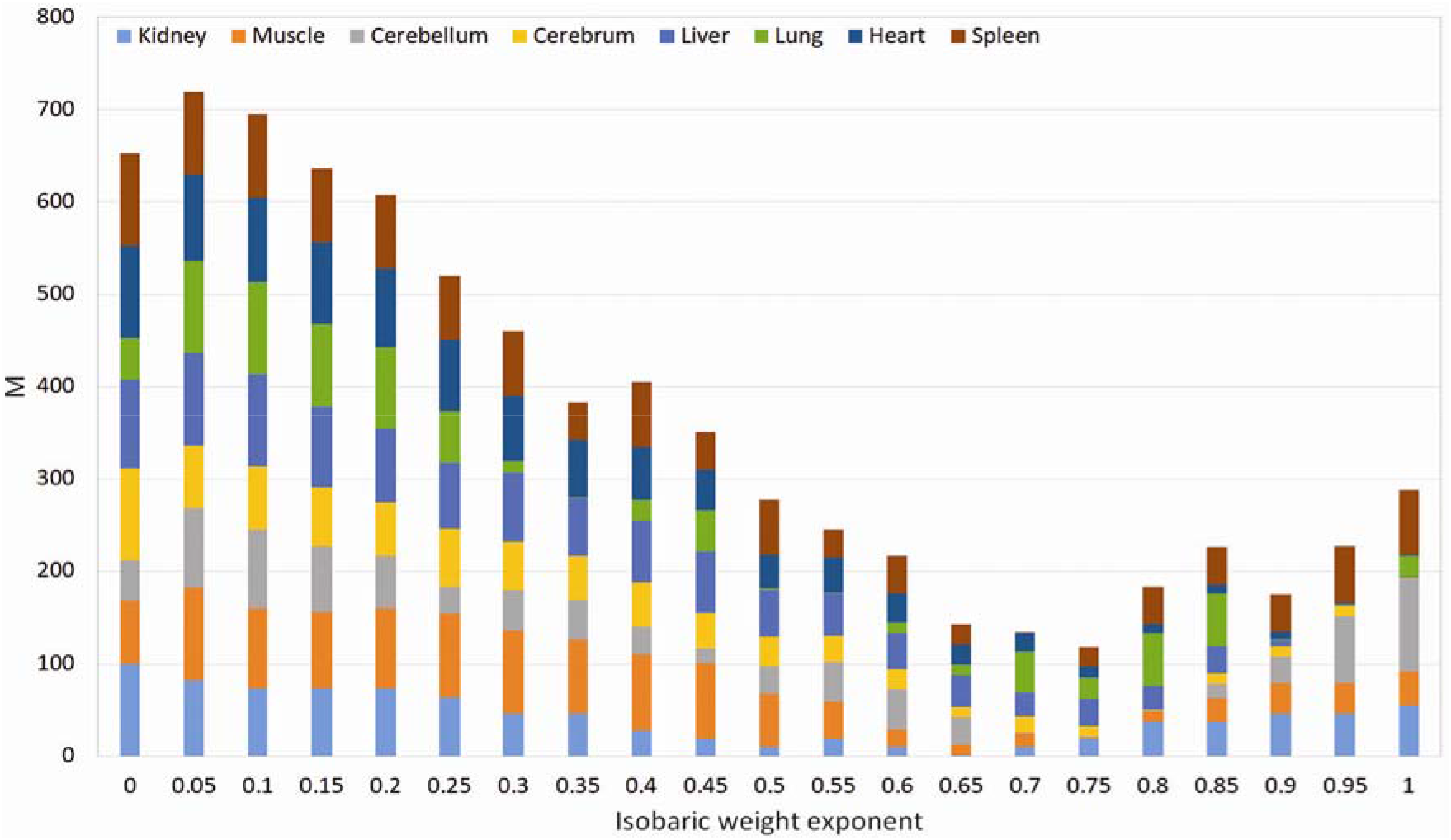
Optimization of the isobaric weight exponent parameter on the dataset by Baileys et al. *M_w_* as defined in the Results section is plotted as a function of the weight exponent. The contributions of separate tissues, *M_g,w_* are color-coded.

### Effects of weighted median-based normalization

In order to test the effect of the novel PSM-level normalization, we ran MaxQuant with and without applying the normalization on the dataset of Bailey *et al*. We then performed UMAP analysis on both outputs. Without normalization, the result of UMAP analysis is dominated by the split into two clusters which separate the data by the acquisition method (Figure 4A). When applying weighted median-based normalization (Figure 4B) UMAP analysis results in strongly focused clusters by tissue across the acquisition methods and all isobaric labeling batches. The impact of the normalization can also be seen by comparing within tissue group variances divided by total variances before and after normalization. Figure 4C shows the media of these over the protein groups. There is a strong decrease in the within group variance brought about by the normalization.

**Figure 4.**
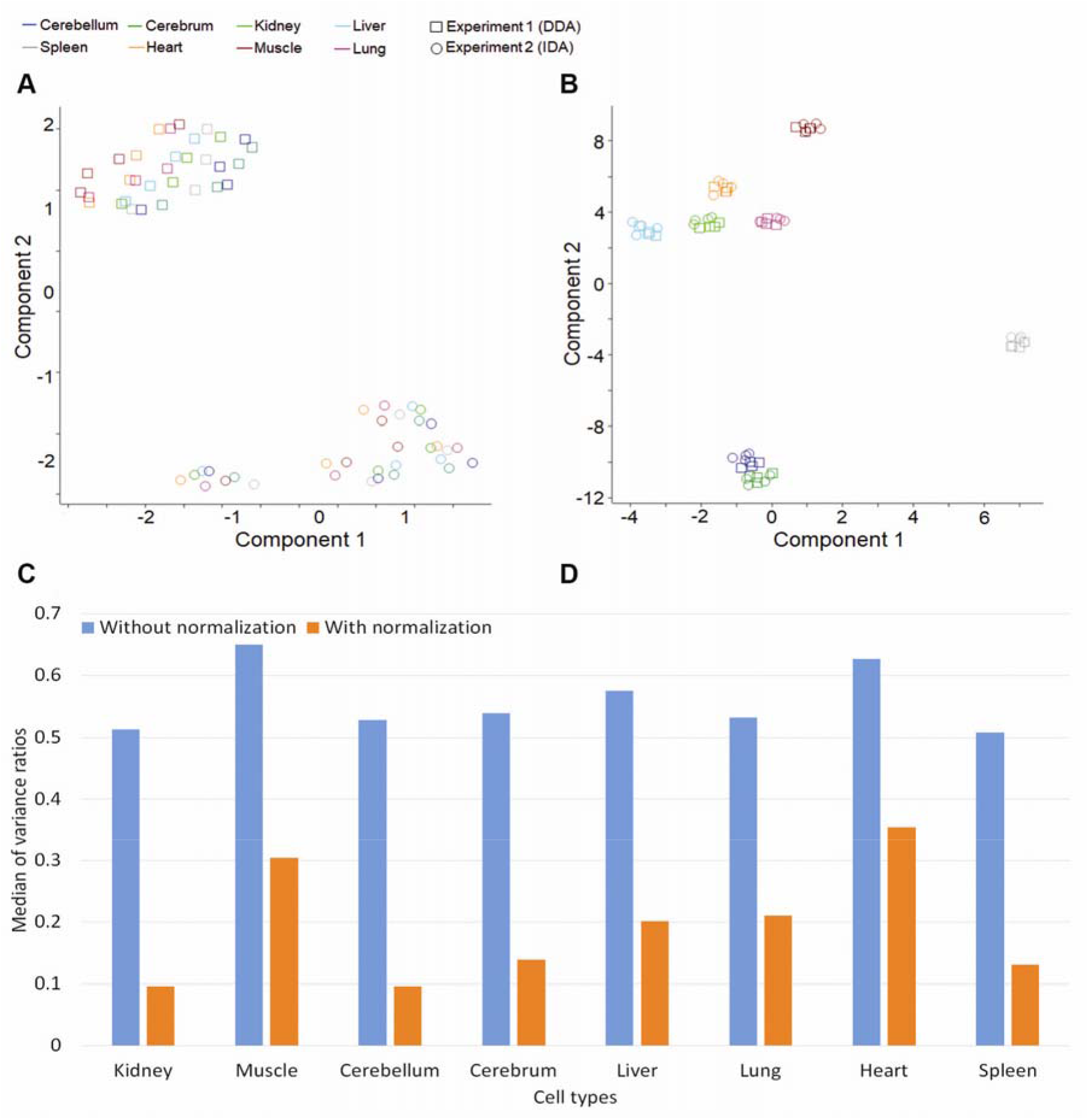
Effect of weighted median normalization. **A.** Result of UMAP analysis on the dataset from Bailey et al. analyzed with MaxQuant without normalization. The different colors designate tissue types while the squares and circles denote the acquisition method. **B.** Same as A., but after applying normalization. **C.** Each bar represents the median over all protein groups of the ratio of within tissue group variances to total variances. Blue and orange bars stand for before and after normalization.

When performing UMAP analysis to the acquisition modes (IDA and DDA) separately, the clustering is also not by tissue but by the TMT 8-plexes (Figure 5A and C). The grouping into TMT-multiplexes coincides with the individual mice, but we assume here that the separation is due to the labeling multiplexes. Independent of what causes the separation, using weighted median normalization removes this batch effect and results in clustering by tissue for both acquisition methods (Figure 5B and D).

**Figure 5.**
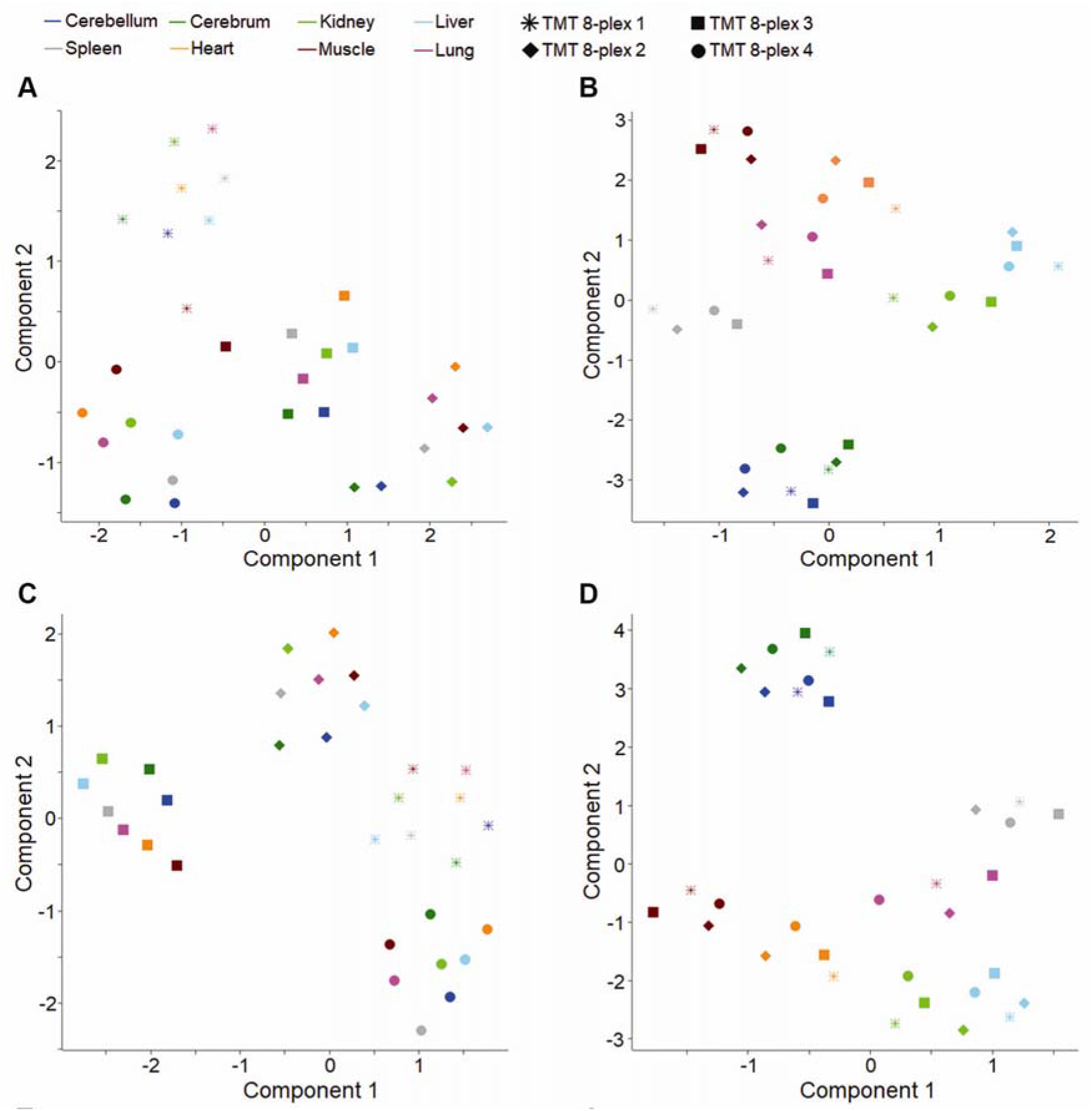
Separate analysis of acquisition methods. **A.** Separate UMAP analysis of the IDA data without normalization Colors represent different tissues, while the symbol types represent different TMT 8-plexes. The samples cluster by 8-plex. **B.** Same as **A.** but with normalization. The samples cluster by tissue. **C.** Same as A for DDA, **D.** Sane as B. for DDA.

### Normalization with reference channels

Here we show the benefits of weighted median normalization for data with reference channels. Similar to the previous section, we perform MaxQuant analysis once without and once with normalization. In both cases, we remove the reference channel and perform UMAP analysis. Without applying normalization, the samples are grouped based on the TMT 10-plex in the UMAP plot (Figure 6A). Using weighted median normalization produces two clusters mainly based on the type of mouse (Figure 6B). Most of the data points not following the clustering by mouse type are the samples from 0 days post infection (dpi). It may be due to the fact that the infection is not active yet at 0 dpi. Hence, we also performed the UMAP analyses of the subset excluding the samples of 0 dpi (Figure 6C). The samples are separated according to WT and *Peli1* knock-out except for one data point. Performing UMAP analysis only for the samples from 20 dpi, the normalized data are completely classified by the type of mice (Figure 6D).

**Figure 6.**
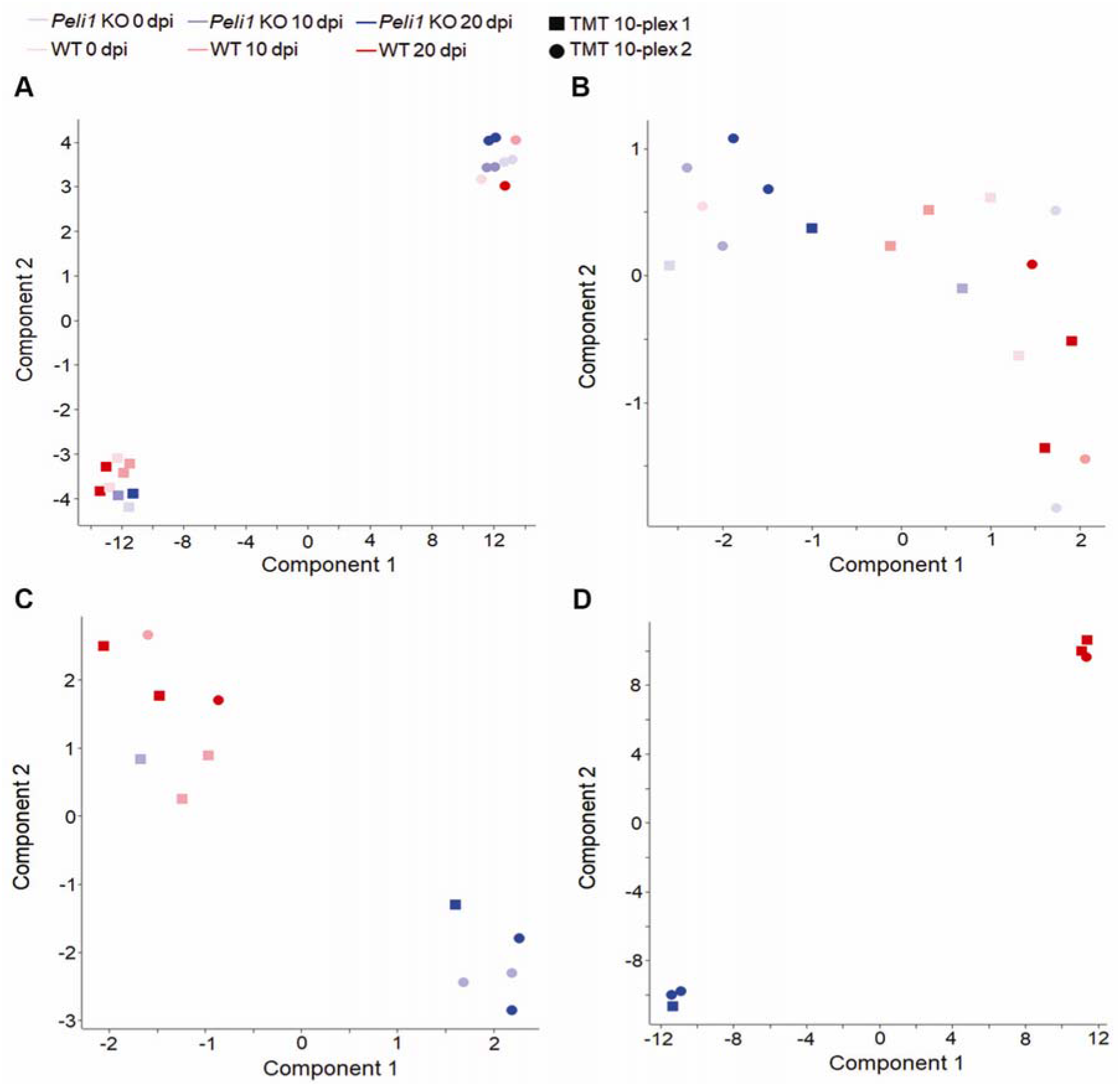
Normalization with reference channel. **A.** Result of UMAP analysis for the dataset by Lereim et al. analyzed with MaxQuant without normalization. The different colors designate conditions and time points while the symbol shapes indicate the TMT batches. **B.** Same as A. but with normalization. **C.** Same as B. but omitting time point 0. **D.** Same as B. but using only the last time point.

### MaxQuant and Perseus plug-ins for isobaric labeling

All of the developed features for isobaric labeling data analysis were integrate into MaxQuant version 1.6.12.0. Reference channels, normalization method and the exponent for the weights of the PSM level ratio normalization can be assigned in the graphical user interface (Figure S1). In order to perform the downstream analysis for the isobaric labeling proteomics data, some newly created plug-ins were integrated in the current version of Perseus (1.6.12.0). All the plug-in activities for isobaric labeling are listed under the heading ‘Isobaric labeling’ in Perseus. These activities are: ‘Annotation from profile’ which can group the samples in isobaric labeling n-plexes based on the profile contained in user defined categorical rows (Table S1 and S2), ‘Remove channels’ for removing the specific channels, for instance those that have been used as a common reference or carrier channel. Moreover, UMAP and t-SNE are also integrated into Perseus (located in ‘Clustering’) for dimension reduction and classification by using PluginInterop and PerseusR^35–38^. The options of the plug-ins can be found in Figure S3 and S4.

## ■ CONCLUSIONS

Two novel approaches for isobaric labeling data analysis were presented, that were integrated into MaxQuant: isobaric match between runs and PSM-level weighted median ratio normalization. They achieve higher precision and fewer missing values in protein quantification. In addition, a collection of Perseus plugins, useful for the downstream analysis of MaxQuant output for isobaric labeling data was introduced. PSM-level normalization efficiently removes batch effects for the subsequent analysis. It is a particularly flexible method, since it does not require to specify what the factor(s) of interest in the dataset are, but works in an unsupervised manner in that respect. Based on results shown in this study, the combination of tools and algorithms in MaxQuant and Perseus is useful gear for the analysis of isobaric labeling MS data.

An alternative method for recovering more MS/MS spectra for TMT quantification beyond the primarily identified PSMs was presented in the context of the single-cell proteomics technology SCoPE-MS^39^. In this complementary approach, the gain in PSMs was achieved by a Bayesian update of the posterior error probability of low-confidence identifications. The combination of this approach with ours could be of interest and will be the subject of further investigations.

Although isobaric labeling with applying IMBR can decrease the number of missing values significantly, not all of the missing values will be replaced with numbers. Hence, imputation is still an important issue that needs to be addressed. Many data analysis methods require a complete data matrix or show potential benefits from imputation. Numerous studies and tools for optimizing imputation have been published and released^12,40,41^. For the remaining missing values in isobaric labeling we propose the same treatment as we recommend for label-free quantification data, which is imputation by drawing from a left-shifted random number distribution, as is done by default in Perseus.

## ■ ASSOCIATED CONTENT

### Supporting Information

- Figure S1: the newly developed features and their setting for isobaric labeling in MaxQuant.
- Figure S2: the workflow in Perseus.
- Figure S3: the plugins for tSNE and UMAP in Perseus.
- Figure S4: the new Perseus plugins of isobaric labeling.
- Table S1: the annotation file for the dataset from Bailey *et al*.
- Table S2: the annotation file for the dataset from Lereim *et al*.

## AUTHOR INFORMATION

### Corresponding Author

*

### Author contributions

S.-Y.H. and J.C. planned and performed the research, developed the software, and wrote the manuscript. S.-Y.H., P.K. and J.C. performed the data analysis.

### Notes

The authors declare no competing financial interest.

## ■ ACKNOWLEDGEMENTS

We thank all members of the Computational Systems Biochemistry research group for helpful discussions.

