## Supplementary material for "Isobaric matching between runs and novel PSM-level normalization in MaxQuant strongly improve reporter ion-based quantification"

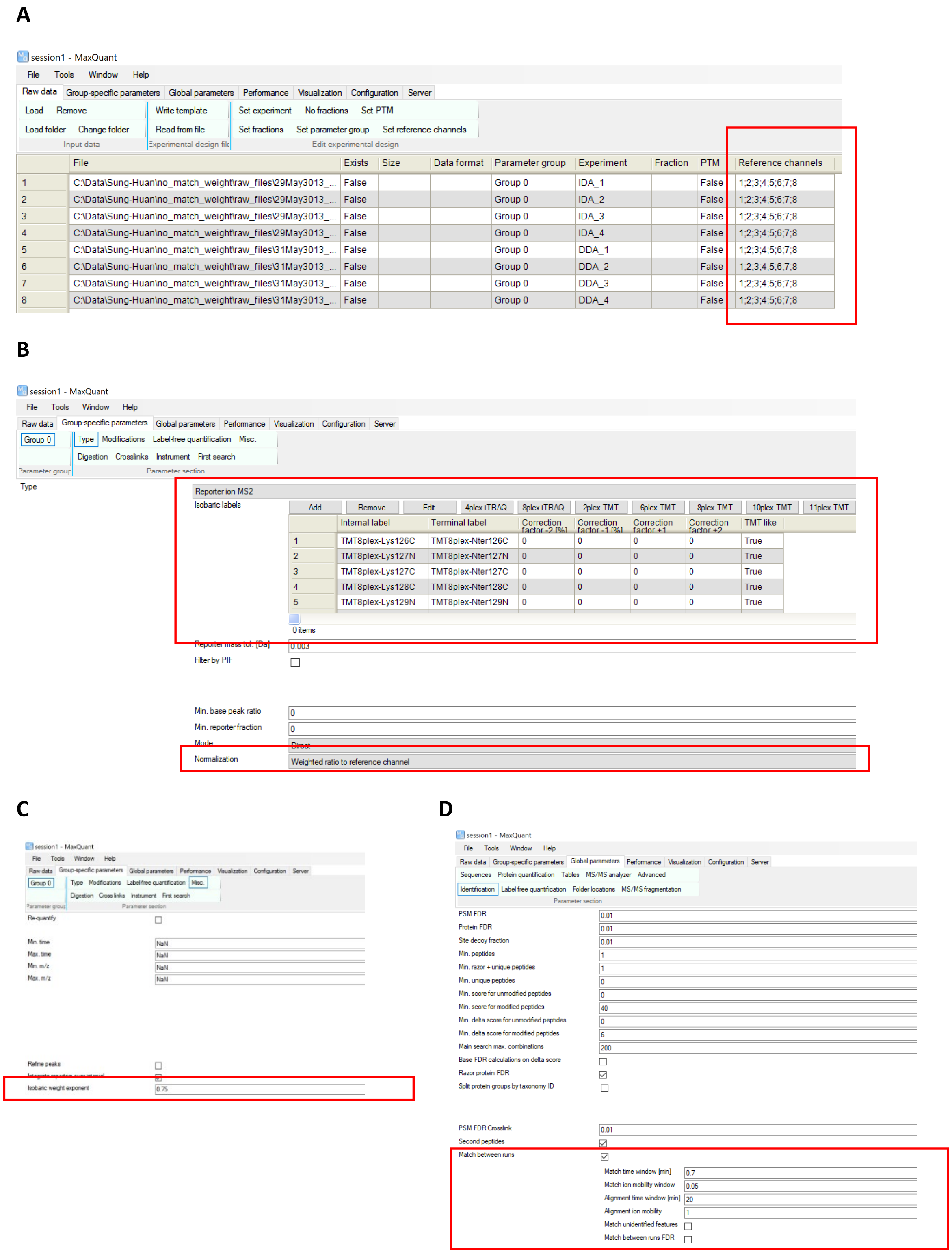


**Figure S1**: The setting of MaxQuant for isobaric labeling proteomics data. **A** The assignment of reference channels. **B** The options for isobaric labeling and normalization method. **C** Isobaric weight exponent. **D** If isobaric labeling is selected in **B**, match between runs will be automatically performed in MS2 level.

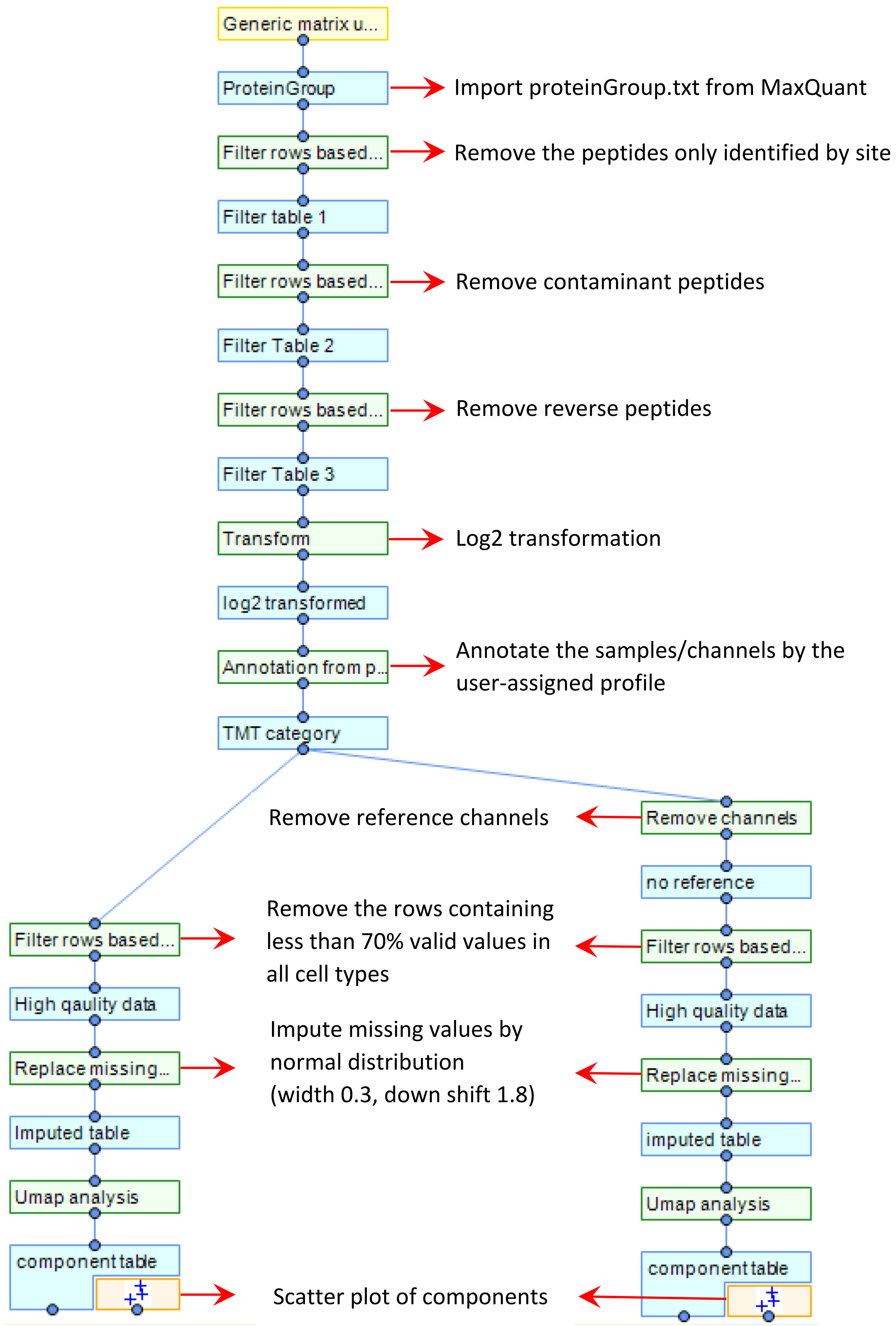


**Figure S2**: Screenshot of a workflow in Perseus. Right branch and left branch are for the dataset with and without reference channels, respectively.

**A**

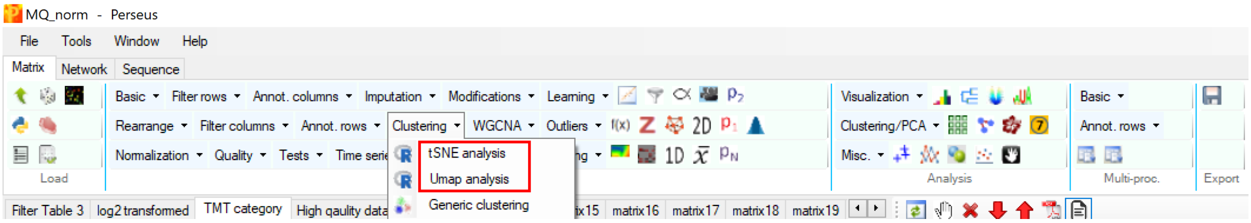


**B**

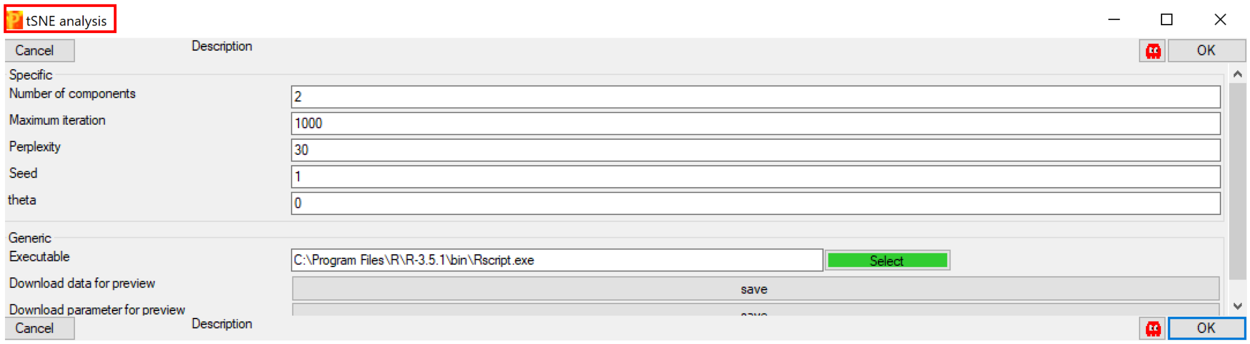


**C**

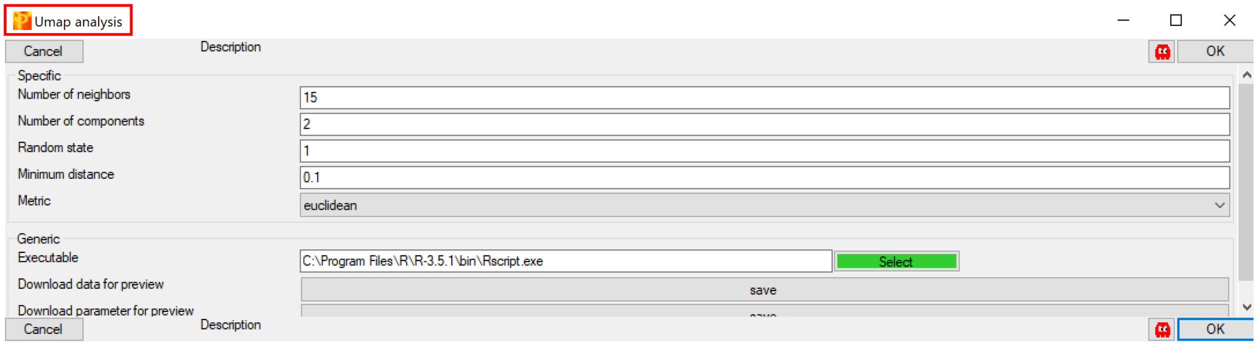


**Figure S3**: Dimensionality reduction methods in Perseus.

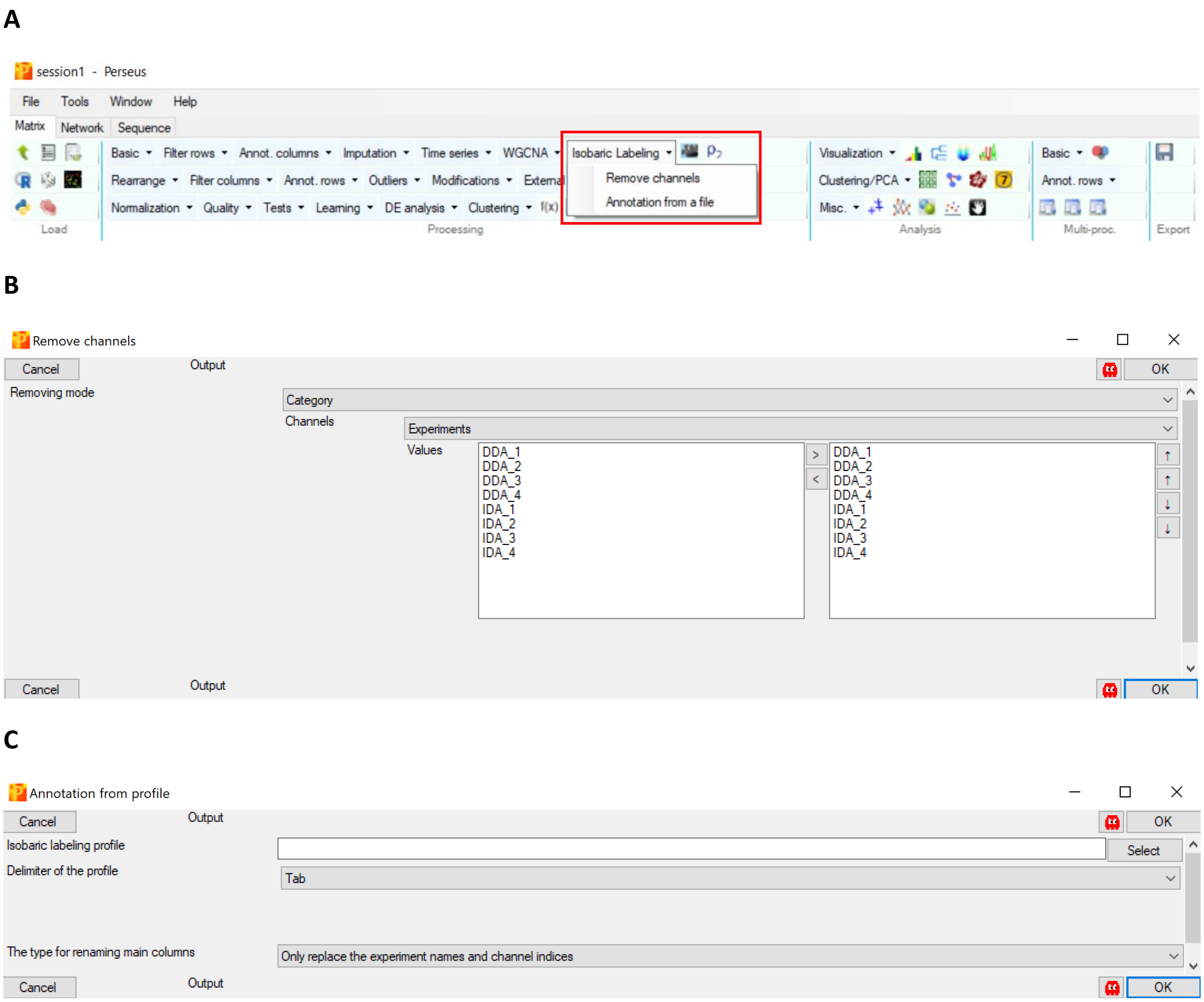


**Figure S4**: The plug-ins for isobaric labeling data in Perseus.

**Table S1**: The annotation profile of Bailey's dataset for "Isobaric Labeling -> Annotation from profile" in Perseus.

**Table S2**: The annotation profile of Lereim's dataset for "Isobaric Labeling -> Annotation from profile" in Perseus.
